# The physiological regulation of macropinocytosis during *Dictyostelium* growth and development

**DOI:** 10.1101/227983

**Authors:** Thomas D. Williams, Robert R. Kay

**Affiliations:** MRC-Laboratory of Molecular Biology, Francis Crick Avenue, Cambridge,CB2 0QH, UK

**Keywords:** *Dictyostelium*, macropinocytosis, endocytosis, flow cytometry

## Abstract

Macropinocytosis is a conserved endocytic process used by *Dictyostelium* amoebae for feeding on liquid medium. To further *Dictyostelium* as a model for macropinocytosis, we developed a high-throughput flow cytometry assay for macropinocytosis, and used it to identify inhibitors and investigate the physiological regulation of macropinocytosis. *Dictyostelium* has two feeding states: phagocytic and macropinocytic. When cells are switched from phagocytic growth on bacteria to liquid media, the rate of macropinocytosis slowly increases, due to increased size and frequency of macropinosomes. Upregulation is triggered by a minimal medium of 3 amino acids plus glucose and likely depends on macropinocytosis itself. Bacteria suppress macropinocytosis while their product, folate, partially suppresses upregulation of macropinocytosis. Starvation, which initiates development, does not of itself suppress macropinocytosis: this can continue in isolated cells, but is shut down by a conditioned-medium factor or activation of PKA signalling. Thus macropinocytosis is a facultative ability of *Dictyostelium* cells, regulated by environmental conditions that are identified here.

**Summary:** A high-throughput flow cytometry assay shows that macropinocytosis in *D. discoideum* is upregulated in the presence of nutrients and absence of bacteria. Development and bacteria induce cells to downregulate macropinocytosis.

## Introduction

Macropinocytosis, first described in the 1930s (Lewis, 1931), is a process of large-scale, non-specific fluid uptake carried out by a wide variety of cells. Actin-driven protrusions from the plasma membrane form cup-shaped circular ruffles that can be several microns in diameter. When a ruffle closes it engulfs and delivers extracellular material to the cell interior in macropinosomes. Macropinosomes proceed through the endocytic system where their contents can be broken down by digestive enzymes and useful metabolites extracted (Buckley and King, 2017).

In the immune system, dendritic cells and macrophages use macropinocytosis to sample environmental antigens for presentation to B and T cells (Sallusto et al., 1995, Norbury et al., 1995). Certain bacteria and viruses can utilise macropinocytosis to invade host cells (Marechal et al., 2001, Nanbo et al., 2010, Hardt et al., 1998), while other bacteria stimulate macropinocytosis to promote toxin internalisation (Lukyanenko et al., 2011). Prions and neurodegenerative protein deposits also invade new host cells through macropinocytosis (Magzoub et al., 2006, Fevrier et al., 2004, Munch et al., 2011, Falcon et al., 2015). Tumour cells can maintain a high rate of macropinocytosis (Lewis, 1937), with Ras-activated cancer cells obtaining a substantial part of their nutrition in this way (Commisso et al., 2013).

Considering its widespread importance, the basic biology of macropinocytosis is poorly understood. It has been studied most intensively in tissue culture cells, particularly macrophages, although genetic screens have been performed in *C. elegans* (Fares and Greenwald, 2001) and *Dictyostelium discoideum* (Bacon et al.,1994). *Dictyostelium* in particular has great potential as a model because of the high constitutive rate of macropinocytosis maintained by cells in the right circumstances and because the evolutionary distance from mammalian cells should allow conserved, core features to be discerned.

The high rate of macropinocytosis by standard axenic strains of *Dictyostelium* used in the laboratory is due to deletion of the RasGAP, NF1 (Bloomfield et al., 2015). This mutation allows cells to grow in nutrient media without a bacterial food source (hence axenic). Wild isolates also perform macropinocytosis, although the rate of fluid uptake is too low to allow growth in the standard media for laboratory-adapted axenic strains. These strains can, however, grow in medium supplemented with additional nutrients (Maeda, 1983, Bloomfield et al., 2015).

Axenic strains form frequent, large macropinosomes. The macropinocytic cups are organized around intense patches of active Ras, Rac and plasmanylinositol (3,4,5)-trisphosphate (PIP3) (Hoeller et al., 2013, Parent et al., 1998, Veltman et al., 2016) (In *Dictyostelium* PIP3 is a plasmanylinositide, rather than a phosphatidylinositide (Clark et al., 2014)), with SCAR/WAVE and WASP localised to their periphery (Veltman et al., 2016). SCAR/WAVE and WASP activate the Arp2/3 complex to polymerise actin and form a macropinocytic cup, which is also known as a crown, or circular ruffle. The cup may be supported by actin polymerization around the base driven by a Ras-activated formin (Junemann et al., 2016).

The rate of fluid uptake through macropinocytosis by axenic cells is regulated by environmental factors, principally whether the cells’ nutrient source is growth media or bacteria (Kayman and Clarke, 1983, Aguado-Velasco and Bretscher, 1999) and their developmental state (Maeda, 1983, Katoh et al., 2007). Macropinocytosis is additionally affected by the stage of the cell cycle and the concentration of bacterial peptone in the medium (Maeda, 1988), as well as the incubation temperature and the pH (Maeda and Kawamoto, 1986). For certain mutants, fluid uptake is dependent upon whether cells are attached to a surface or in shaking suspension (Novak et al., 1995).

Fluid uptake by standard axenic strains of *Dictyostelium*, such as Ax2, is almost entirely due to macropinocytosis and can be accurately measured by following the uptake of fluorescent dextran as a fluid phase marker (Kayman and Clarke,1983, Thilo and Vogel, 1980, Hacker et al., 1997). We have developed a high-throughput assay to measure macropinocytosis in *Dictyostelium*, identified useful inhibitors and sought to better understand how macropinocytosis is physiologically regulated during the switch between macropinocytic and phagocytic feeding and the growth-to-development transition.

## Results

### Measurement of uptake by high-throughput flow cytometry

Macropinocytosis accounts for more than 90% of fluid uptake by axenic strains of *Dictyostelium*, and can therefore be followed by measuring fluid uptake (Hacker et al., 1997). However, existing methods based on processing individual cell pellets after uptake of fluorescent dextran are of relatively low throughput. We therefore developed a high-throughput assay using flow cytometry to measure TRITC-dextran uptake. The assay is performed in 96-well plates and, after loading with TRITC-dextran, the cells are washed *in*-*situ* by ‘dunk-banging’ and detached using sodium azide (Glynn and Clarke, 1984) (Figure 1A), which also prevents exocytosis of internalized dextran (Figure 1B). Plates are analysed by flow cytometry using a High-Throughput Sampling attachment to load the flow cytometer, and subsequent analysis is performed using Flowjo, which easily distinguishes *Dictyostelium* cells from beads and bacteria, but not yeast (Figure 1C). A principle advantage of flow cytometry is that the fluorescence of internalized TRITC-dextran can be determined for single cells (Figure 1D). The accumulation of TRITC-dextran proceeds in a uniform fashion across the population over time, with an extended lagging edge of cells with lower uptake. Median fluid internalisation over time by Ax2 cells is quantified in figure 1E.

**Figure 1.**
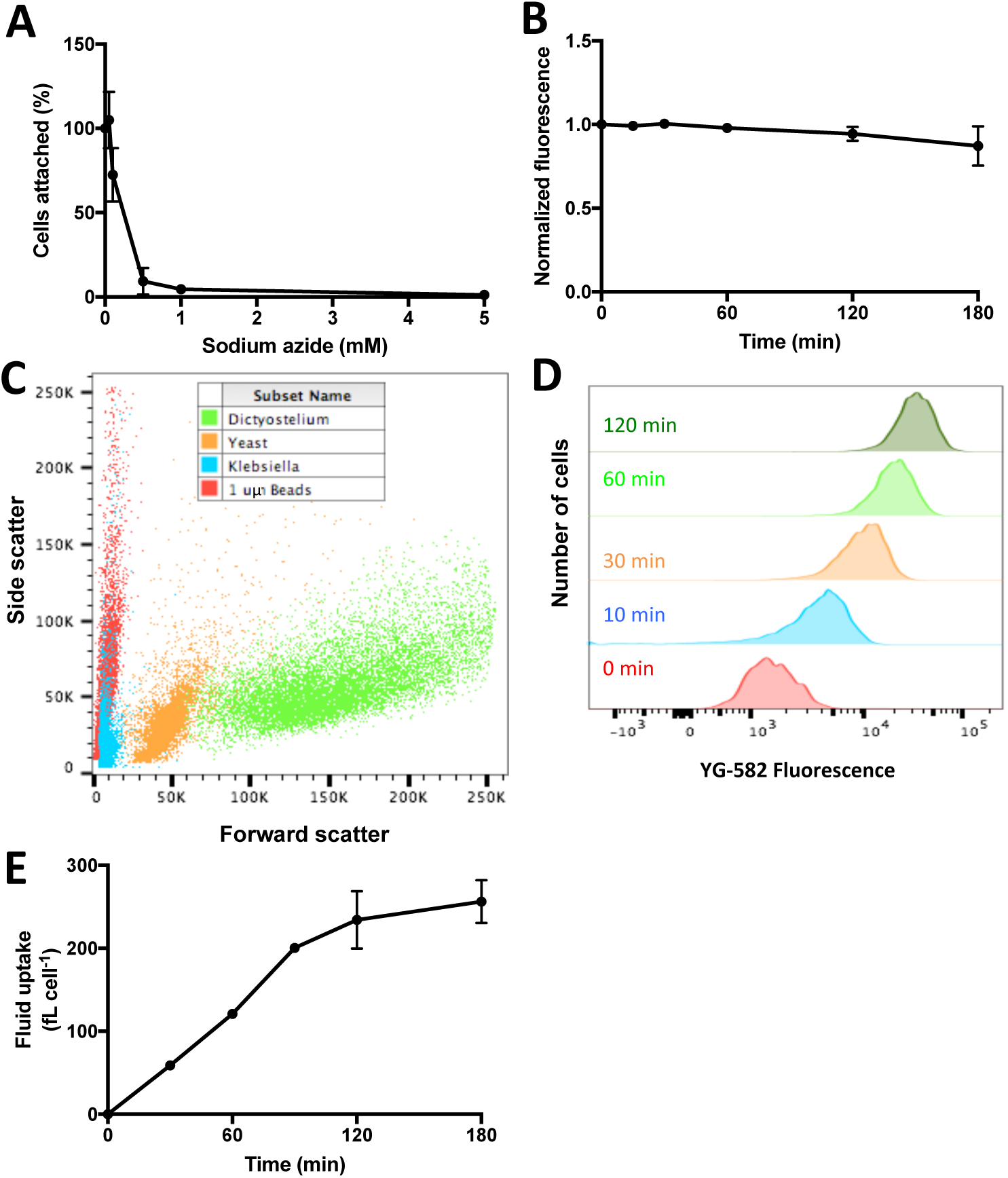
Fluid uptake measurement by high-throughput flow cytometry. **A)** Sodium azide causes efficient detachment of cells in 96-well plates. Attached cells were incubated with sodium azide for 5 minutes and the proportion remaining attached was measured using crystal violet staining (Bloomfield et al., 2015). **B)** Sodium azide prevents significant exocytosis of TRITC-dextran for at least 2-3 hours. Cells, loaded with dextran, were washed and incubated in 5 mM sodium azide and intracellular fluorescence measured by flow cytometry. **C)** Representative dot-plots showing forward and side scatter for beads, bacteria, yeast and *Dictyostelium* cells. *Dictyostelium* is easily distinguished from bacteria, beads and background particles by gating, but cannot be separated fully from yeast particles. **D)** Representative histograms showing the internalised TRITC-dextran of individual cells within a population over time. Axenically grown Ax2 cells were incubated in shaking suspension with TRITC-dextran for up to two hours and analysed by flow cytometry. TRITC-dextran accumulates in every cell, although there is a lagging tail of cells with lower fluid uptake. **E)** Fluid uptake time-course of Ax2 cells in a 96-well plate. TRITC-dextran uptake proceeds linearly for the first 60-90 minutes, then plateaus as it begins to be exocytosed. All error bars show s.e.m.; n=3 in all experiments.

Controls show the efficiency of the wash step (Figure S1A); that Ax2 cells take up similar volumes of liquid whether in suspension or attached to a surface (as in the assay-Figure S1B), although this is not true for all strains (Novak et al.,1995); and the range of cell numbers that can be accommodated per well (Figure S1C). The assay is calibrated in terms of volume taken up per cell by reference to measurements of uptake by the same cell population using a fluorimeter (Figure S1D) and standardised over time using Flow-Set fluorosphere calibration beads (Beckman Coulter). Phagocytosis of beads (Figure S1E) and bacteria (Figure S1F) as well as membrane uptake (Figure S1G) can be readily measured.

### Effect of inhibitors on macropinocytosis

Inhibitors are powerful tools for acutely interfering with biological processes, but relatively few are currently known for macropinocytosis. We therefore tested a number of inhibitors affecting both the cytoskeleton and cellular signalling. These were added to Ax2 cells growing in HL5 medium at the start of the uptake assay, with the TRITC-dextran. The internalised fluorescence was measured 1 hour later (Table 1). A number of inhibitors were without effect, although whether this is due to lack of inhibitor uptake, target interaction or the target not functioning in macropinocytosis is unknown.

**Table 1.** Effect of inhibitors on macropinocytosis. Inhibitors were added at several concentrations to axenically growing Ax2 cells in 96-well plates in conjunction with TRITC-dextran for one hour and the fluid uptake of the cells during that time determined. Dose response curves are shown for the inhibitors that were effective in inhibiting macropinocytosis in figure S2. These were repeated three times.

| Inhibitor | Source | Molecular target(s) | Inhibits macropinocytosis? | IC<sub>50</sub> (μM) | Maximum inhibition (μM) | Maximum dose tested (μM) |
| --- | --- | --- | --- | --- | --- | --- |
| Latrunculin B | Sigma-Aldrich | F-actin | Yes | 1 | 5 | 10 |
| Cytochalasin A | Cayman | F-actin | Yes | 1.7 | 10 | 10 |
| Cytochalasin B | Sigma-Aldrich | F-actin | No | N/A | N/A | 200 |
| Cytochalasin D | Cayman | F-actin | No | N/A | N/A | 200 |
| Jasplakinolide | Santa Cruz | F-actin | No | N/A | N/A | 20 |
| CK666 | Sigma-Aldrich | Arp2/3 | Yes | 30 | 60 | 100 |
| SMIFH2 | Sigma-Aldrich | Formins | Yes | 5 | 30 | 100 |
| Wiskostatin | Sigma-Aldrich | WASP | Yes | 2.75 | 10 | 10 |
| Nocodazole | Sigma-Aldrich | Microtubules | Yes | 70 | 70 | 300 |
| Thiabendazole | Sigma-Aldrich | Microtubules | Yes | 70 | 150 | 200 |
| Blebbistatin | Sigma-Aldrich | Myosin II | No | N/A | N/A | 100 |
| Dynasore | Sigma-Aldrich | Dynamin | No | N/A | N/A | 310 |
| LY294002 | Cayman | PI3K, TORC2 | Yes | 38 | 75 | 200 |
| BYL719 | Cayman | PI3K alpha | No | N/A | N/A | 250 |
| TGX221 | Cayman | PI3K beta | Yes | 60 | 150 | 200 |
| CAL101 | Cayman | PI3K gamma | No | N/A | N/A | 250 |
| EHT1864 | Cayman | Rac | Yes | 0.03 | 0.1 | 0.1 |
| Rapamycin | Sigma-Aldrich | TORC1 | No | N/A | N/A | 10 |
| PP242 | Sigma-Aldrich | TORC1, TORC2 | No | N/A | N/A | 200 |
| Palomid 529 | Sigma-Aldrich | TORC1, TORC2 | No | N/A | N/A | 500 |
| Torin 1 | Sigma-Aldrich | TORC1, TORC2 | Yes | 20 | 50 | 125 |
| Amiloride | Adooq Biosciences | Na<sup>+</sup>/H<sup>+</sup> exchanger | No | N/A | N/A | 200 |
| EIPA | Cayman | Na<sup>+</sup>/H<sup>+</sup> exchanger | No | N/A | N/A | 200 |
| EGTA | Sigma-Aldrich | Extracellular calcium | No | N/A | N/A | 2000 |

As macropinocytosis is an actin-dependent process, we tested a number of inhibitors of actin dynamics. Latrunculin B efficiently inhibited macropinocytosis at standard concentrations (Figure S2A), as expected from its profound effects on the actin cytoskeleton (Pramanik et al., 2009). Cytochalasin A, as previously reported (Hacker et al., 1997), inhibited macropinocytosis (Figure S2B). Inhibitors of the Arp2/3 complex (CK666, Figure S2C), WASP (Wiskostatin, Figure S2D) and formins (SMIFH2, Figure S2E) were all potent inhibitors of macropinocytosis, consistent with the localization of the target proteins to macropinosomes, the macropinocytosis defects in WASP (Veltman et al., 2016) and ForG (Junemann et al., 2016) mutants and the axenic growth defect of an ArpB mutant (Langridge and Kay, 2007).

The microtubule inhibitors nocodazole (Figure S2F) and thiabendazole (Figure S2G) both partially inhibited fluid uptake, indicating a role for microtubules. The myosin II inhibitor blebbistatin had no effect on macropinocytosis, in contrast to previously published data (Shu et al., 2005). This may be because, in our hands, blebbistatin readily precipitated at concentrations above those used.

Macropinosomes are organised around active Rac/Ras/PIP3 patches (Hoeller et al., 2013, Veltman et al., 2016) and, accordingly, the PI3-kinase (PI3K) inhibitor LY294002 (Figure S2H) inhibits fluid uptake. TGX221, which targets the mammalian p110β PI3K isoform, inhibited macropinocytosis (Figure S2I) whereas inhibitors targeting α and γ isoforms did not. We found the Rac inhibitor EHT1864 (Shutes et al., 2007) is a potent inhibitor of fluid uptake (Figure S2J).

In mammalian cells macropinocytosis of free amino acids, notably leucine, induces mTORC1 activation (Yoshida et al., 2015), allowing cell proliferation. Rapamycin, a TORC1 specific inhibitor, did not affect fluid uptake when applied acutely, as found previously, although it does prevent proliferation (Rosel et al., 2012). It has been suggested that there are functions of mTORC1 that are not inhibited by rapamycin, but are by more potent, less specific, mTor inhibitors (Thoreen and Sabatini, 2009). We therefore tried alternative Tor inhibitors and observed an inhibition of macropinocytosis in cells treated with torin 1 (Figure S2K), but not palomid 529 or PP242. Whether this is due to greater inhibition of TORC1, inhibition of TORC2, or both is not clear: TORC2 has previously been described as having no function in *Dictyostelium* macropinocytosis (Rosel et al., 2012), however we see a reduction in macropinocytosis when TORC2 components are knocked out in the Ax2 strain used here (not shown).

The nearest to diagnostic inhibitors for macropinocytosis in mammalian cells are amiloride and EIPA, which block the plasma membrane Na+/H+ exchanger, thus affecting sub-membranous pH (Koivusalo et al., 2010). Although *Dictyostelium* possesses two Na+/H+ exchangers (Patel and Barber, 2005, Fey et al., 2013), it is not known whether they are sensitive to these drugs and we find that the drugs do not affect macropinocytosis. The removal of extracellular calcium by EGTA inhibits constitutive macropinocytosis in immune cells (Canton et al., 2016), but had no effect on macropinocytosis by *Dictyostelium* incubated in a calcium-free medium (50 mM lysine, 55 mM glucose in 50 mM MES, pH 6.5) indicating that extracellular calcium is not required. Indeed, high extracellular calcium concentrations are reported to inhibit *Dictyostelium* macropinocytosis (Maeda and Kawamoto, 1986).

These results support previous genetic studies showing that macropinocytosis depends on PI3K, Rac and actin dynamics controlled through SCAR/WAVE, WASP and formins. On the other hand, regulation through extracellular calcium is not part of a core, conserved mechanism of macropinocytosis across species, while roles for the Na+/H+ exchanger and Tor have not been confirmed.

### Slow switching between feeding strategies

Ax2 cells grown on bacteria have a low rate of macropinocytosis, which increases greatly when they are switched to HL5 growth medium (a complex medium containing peptone, yeast extract and glucose). A similar, although much reduced, increase is seen in wild-type NC4 cells (which have an intact NF1 gene) in media enriched with protein (Maeda, 1983).

We confirmed the upregulation of macropinocytosis in Ax2 cells switched from growth on bacteria to HL5 (Figure 2A). It is slow, taking about 10 hours (Figure 2B), similar to Ax3 (Kayman and Clarke, 1983), and involves both an increased rate of macropinosome formation (Figure 2C) and size (Figure 2D). A 50% increase in diameter, as seen here, would lead to a 3-4 fold increase in macropinosome volume.

**Figure 2.**
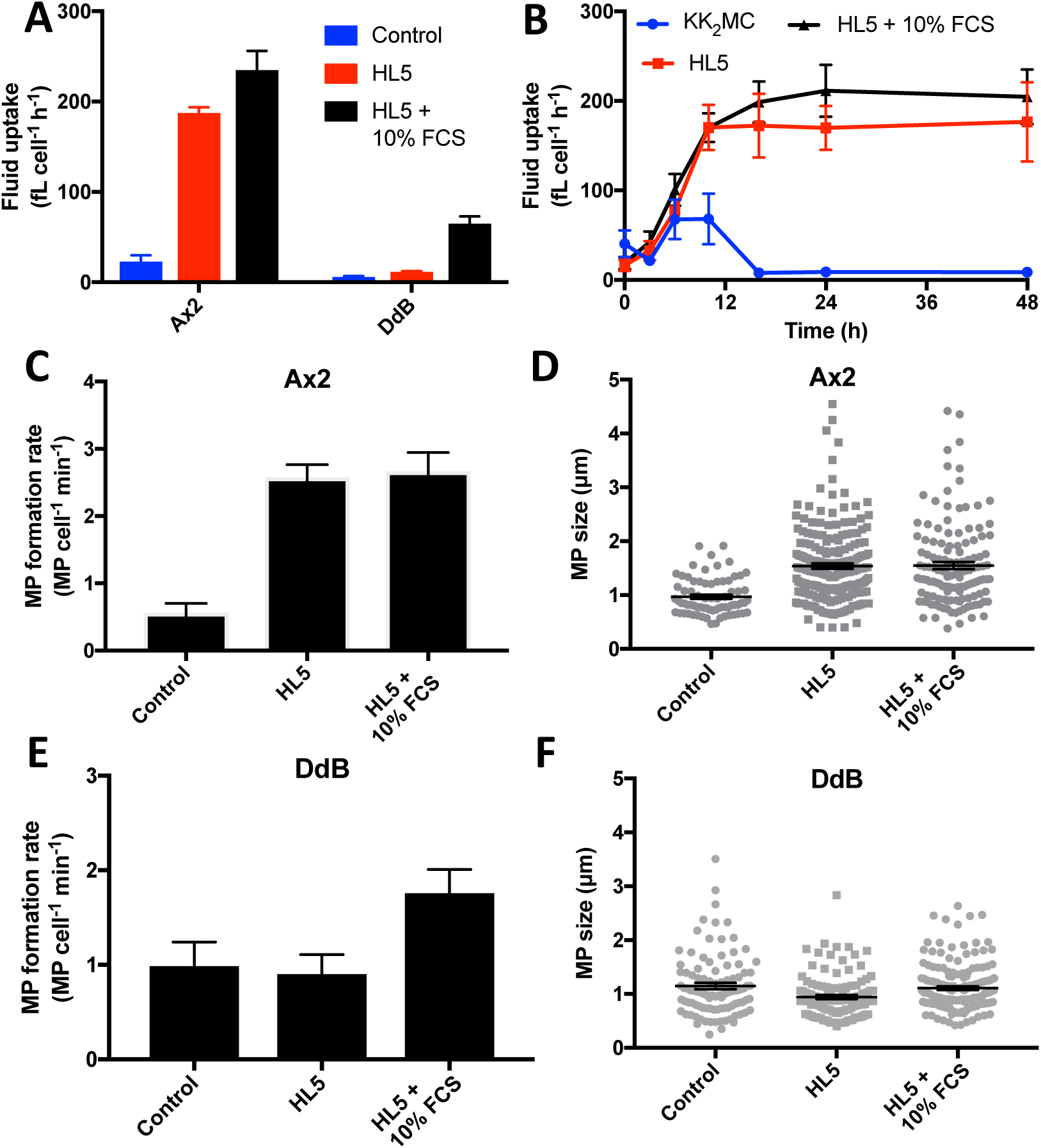
Cells adapt to growth on liquid media by increasing their rates of fluid uptake and macropinocytosis. **A)** Macropinocytosis increases when cells grown on bacteria are transferred to liquid medium. Fluid uptake was either measured immediately after harvesting cells from bacteria (control) or after 24 hours in the indicated media (n=3). **B)** Kinetics of the increase in fluid uptake by Ax2 cells during adaptation to nutrient media (n=3). **C)** The rate of macropinosome formation increases in Ax2 cells adapted to nutrient media. Macropinosome formation was measured microscopically after a 1-minute pulse with FITC-dextran (n=7). **D)** The size of macropinosomes increases in Ax2 cells adapted to nutrient media. The maximum diameter of macropinosomes at the moment of closure was measured in the mid-section of cells using the PIP3 reporter PkgE-PH mCherry on 3 separate days. **E)** Macropinosome formation increases in DdB cells adapted to HL5 fortified with 10% FCS (n=6). **F)** Macropinosome size does not increase in DdB cells adapted to liquid media. Cells were imaged on 3 separate days.Ax2 is a standard laboratory strain able to grow in HL5 medium, which has the NF1 gene, deleted; DdB is its non-axenic parent with an intact NF1 gene. In all experiments cells were grown on bacteria, washed and then transferred to the indicated media. Excluding panel B, the control measurements were made with cells freshly harvested from bacteria and the others after 24 hours incubation in the indicated media. Fluid uptake and other measurements were made as described in the Materials and Methods. Error bars show the s.e.m.

Wild-type DdB cells (the parent of the standard Ax2, Ax3 and Ax4 strains) with an intact NF1 gene do not noticeably upregulate macropinocytosis in HL5 (Figure 2A). However, if DdB cells are switched to HL5 supplemented with 10% FCS (Gibco, providing ~4 mg ml^-1^ additional protein), in which they can proliferate (Bloomfield et al., 2015), they substantially upregulate macropinocytosis, although not as much as Ax2. The increased fluid uptake by DdB cells in this case appears to be due only to an increased rate of macropinosome formation (Figure 2E), with no detectable increase in size (Figure 2F). Thus the macropinocytic rate of wild-type cells is also controlled by the availability of environmental nutrients, as in Ax2 cells.

Ax2 and other axenic cells can consume yeast and other large particles, but this ability depends on the loss of the NF1 gene and is not shared by wild-type cells (Bloomfield et al., 2015). Thus the abilities to phagocytose large particles and to take in large volumes of fluid by macropinocytosis are linked as both depend on the loss of NF1. We asked whether this linkage is also seen at a physiological level. It is apparent from Figure S3A that it is: Ax2 cells adapted to HL5 medium and with a high macropinocytic rate can take up yeast or large beads (2 micron diameter) much better than the same cells grown on bacteria, which have a low macropinocytic rate. Uptake of either particle by DdB cells, which have an intact NF1 gene, was unaltered by the nutritional environment they came from (Figure S3B), consistent with the unaltered macropinosome size observed in figure 2F.

The macropinocytic and phagocytic states are not mutually exclusive, as we found that Ax2 cells fully adapted to HL5 maintain a relatively high rate of phagocytosis of bacteria (Figure S3C). We therefore asked what happens when Ax2 cells are presented with both bacteria and liquid medium for food. In this case, irrespective of whether the cells had been grown on bacteria or HL5, they adopted a low rate of macropinocytosis (Figure S3D).

These results show that *Dictyostelium* has two basic feeding modes: the preferred mode is phagocytosis, which is seen with cells growing on bacteria. Alternatively, cells in liquid media without bacteria adopt a second mode, in which macropinocytosis is upregulated, although the potential for phagocytosis of bacteria is retained.

### A minimal set of soluble nutrients can stimulate the upregulation of macropinocytosis

Ax2 cells do not sustainably increase their rate of macropinocytosis when they are switched from bacteria to buffer (Figure 2B), but require HL5 medium, or some components of it, to do so. To identify such stimulatory components, we first showed that HL5 could be replaced by the SIH defined medium (Figure 3A) and then dissected this defined medium to find the active components. Leaving out blocks of components showed that vitamins and micro-minerals are not necessary for macropinocytic upregulation and that the effect is accounted for by amino acids and glucose alone (Figure 3B). Testing amino acids individually showed that only arginine, lysine and glutamate induce macropinocytosis upregulation at the tested concentrations (Table S1). Consistent with this, removal of these amino acids from SIH severely impairs the ability of cells to upregulate macropinocytosis, which is restored when the amino acids are returned to the medium (Figure 3C). Testing different sugars showed that only glucose and other metabolizable sugars that can support cell growth permit macropinocytosis upregulation (Table S2) (Watts and Ashworth, 1970,Ashworth and Watts, 1970).

**Figure 3.**
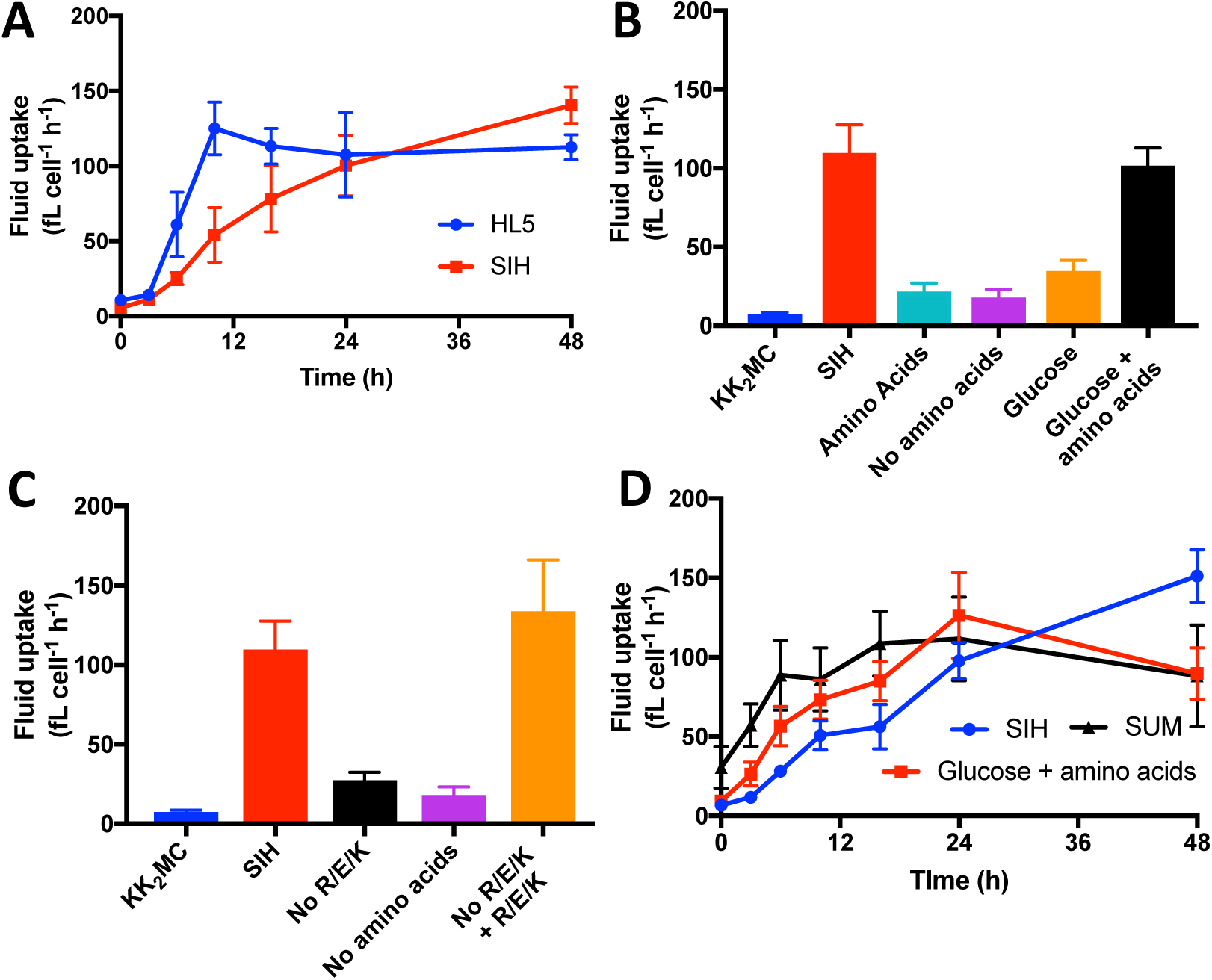
Macropinocytosis upregulation can be induced by a minimal medium containing glucose, arginine, lysine and glutamate. **A)** The defined SIH medium efficiently induces upregulation of macropinocytosis in cells transferred from bacteria. The complex HL5 medium is shown for comparison (n=3). **B)** Broad dissection of SIH medium shows that the amino acids and glucose are responsible for its ability to stimulate macropinocytosis upregulation (n=7). **C)** Detailed dissection of SIH medium (see supplementary tables 1 and 2) shows that arginine, glutamate and lysine (R, E and K) are needed for efficient upregulation of macropinocytosis (n=5). **D)** A minimal medium containing arginine, glutamate, lysine and glucose (SUM) gives efficient upregulation of macropinocytosis. The kinetics of upregulation induced by SUM, glucose and amino acids and SIH are compared (n=3). Ax2 cells grown on bacteria were washed free of bacteria and transferred to the indicated media for 24 hours, unless indicated otherwise, and then fluid uptake measured by flow cytometry as described in materials and methods. Error bars show the s.e.m.

Based on these results, a simplified medium for macropinocytosis upregulation, SUM (Simple Upregulation Media) was devised, consisting of KK_2_MC plus 55 mM glucose, 4 mM arginine, 3.7 mM glutamate and 8.5 mM lysine (the same concentrations as SIH) at pH 6.5. SUM induces nearly the same level of macropinocytosis as complete SIH, with faster upregulation kinetics (Figure 3D). Although cells remain healthy in SUM for several days, it does not support longterm growth. SUM has very low background fluorescence, and we have found it very useful for microscopy, particularly for cells with weakly expressed markers, such as knock-ins. Cells can be grown rapidly on bacteria before transfer to SUM a few hours prior to microscopy, during which time macropinocytosis is greatly upregulated.

These results show that macropinocytosis upregulation can be induced by only a handful of the components present in defined medium, while the requirement for the sugar to be metabolizable hints that sugars may be sensed through their effects on metabolism, rather than by dedicated receptors.

### Macropinocytosis is required for efficient upregulation of macropinocytosis

We envisioned that nutrients that cause macropinpocytosis upregulation might either be sensed by dedicated receptors, such as for glutamate, or indirectly through their effect on metabolism, or a combination of both. Since nutrients obtained by macropinocytosis can only be utilized after internalization and digestion, this second route implies that macropinocytic upregulation would depend on macropinocytosis itself. To test this idea we turned to the inhibitors identified earlier in this work to inhibit macropinocytosis during upregulation.As this experiment requires prolonged inhibitor treatment, we first tested how well cells recover from the inhibitors. Ax2 cells growing in HL5 recover quite well from prolonged treatment with LY294002, TGX221 (both PI3K), CK666 (Arp2/3 complex), EHT1864 (Rac) and torin 1 (Tor) (Figure S4A-E). Prolonged incubation with other inhibitors was too deleterious to make them useful.

We next used the inhibitors to determine to what extent the upregulation of macropinocytosis is affected by inhibiting macropinocytosis (Figure 4 ‘raw’ curves), also making a correction for the relatively small deleterious effects of long-term exposure of cells to the inhibitors (Figure 4, ‘corrected’ curves). Although these inhibitors affect macropinocytosis through different targets, they all inhibit upregulation (measured after 10 hours incubation in HL5) in a dose-dependent manner (Figure 4A-E). The effect remains even after correcting for the long-term effects of the inhibitors. Upregulation is not completely abolished by the inhibitors, reflecting their incomplete inhibition of macropinocytosis.Thus these results suggest that the upregulation of macropinocytosis by nutrient media depends on delivery of nutrients into the cell through macropinocytosis.

**Figure 4.**
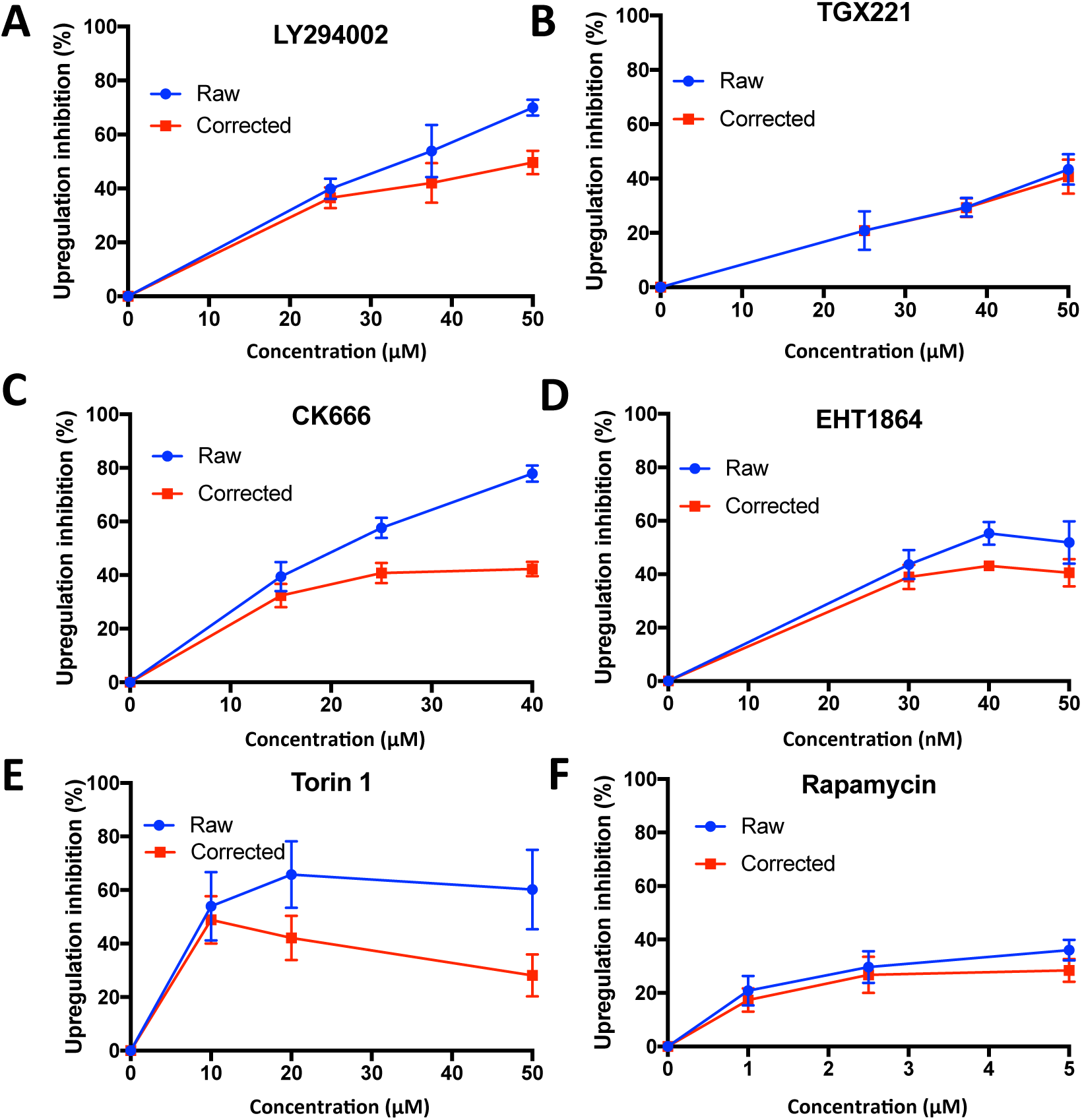
Evidence that macropinocytosis upregulation depends on macropinocytosis. To test whether macropinocytosis upregulation depends on macropinocytosis, inhibitors with differing targets (see Table 1) were used to inhibit macropinocytosis during the upregulation period. The inhibitor was then washed away and the degree of upregulation determined by measuring fluid uptake compared to untreated controls (‘raw’ curves). To control for long-term effects of the inhibitors, cells with fully upregulated macropinocytosis were treated in parallel and the results corrected accordingly (‘corrected’ curves; see Figure S4). Inhibitors used and their nominal targets: **A)** LY29004 (PI3-kinase, n=3); **(B)** TGX221 (PI3-kinase, n=4); **(C)** CK666 (Arp2/3 complex, n=4); **(D)** EHT1864 (Rac, n=4); **(E)** Torin 1 (Tor, n=3). **F)** Rapamycin (TORC1, n=3). Ax2 cells, harvested from bacteria, were incubated in HL5 in 96-well plates with the inhibitors for 10 hours, then the inhibitors washed away by dunk-banging, the cells allowed to recover for 10 minutes and the fluid uptake measured over 1 hour using the high-throughput flow cytometry assay. To correct for deleterious effects of the inhibitors, control Ax2 cells grown in HL5 (with maximally upregulated macropinocytosis) were similarly treated with inhibitors for 10 hours and their fluid uptake compared to untreated controls to give the correction factor: Uptake(drug-treated control cells)/Uptake(vehicle-treated control cells) by which the raw data was multiplied to give the corrected curves. Error bars show the s.e.m.

We considered the possibility that the ingested nutrients delivered by macropinocytosis might be detected through the TORC1 complex, similar to other organisms. Although rapamycin does not inhibit macropinocytosis acutely, it does somewhat inhibit upregulation (Figure 4F), with extremely mild effects on control cells (Figure S4F). Torin1, has a stronger effect on upregulation (Figure 4E), but as it is less specific, some of this might be due to inhibition of the TORC2 complex. In summary, these results suggest that nutrients causing cells to increase their rate of macropinocytosis are detected in the macropinocytic pathway, possibly by TORC1.

### Sensing of bacteria

Bacteria have two distinct effects on the regulation of macropinocytosis. They inhibit upregulation of macropinocytosis by cells transferred to HL5 (Figure 5A) and promote downregulation of macropinocytosis by cells transferred from HL5 to buffer, where it otherwise remains high (Figure 5B).

**Figure 5.**
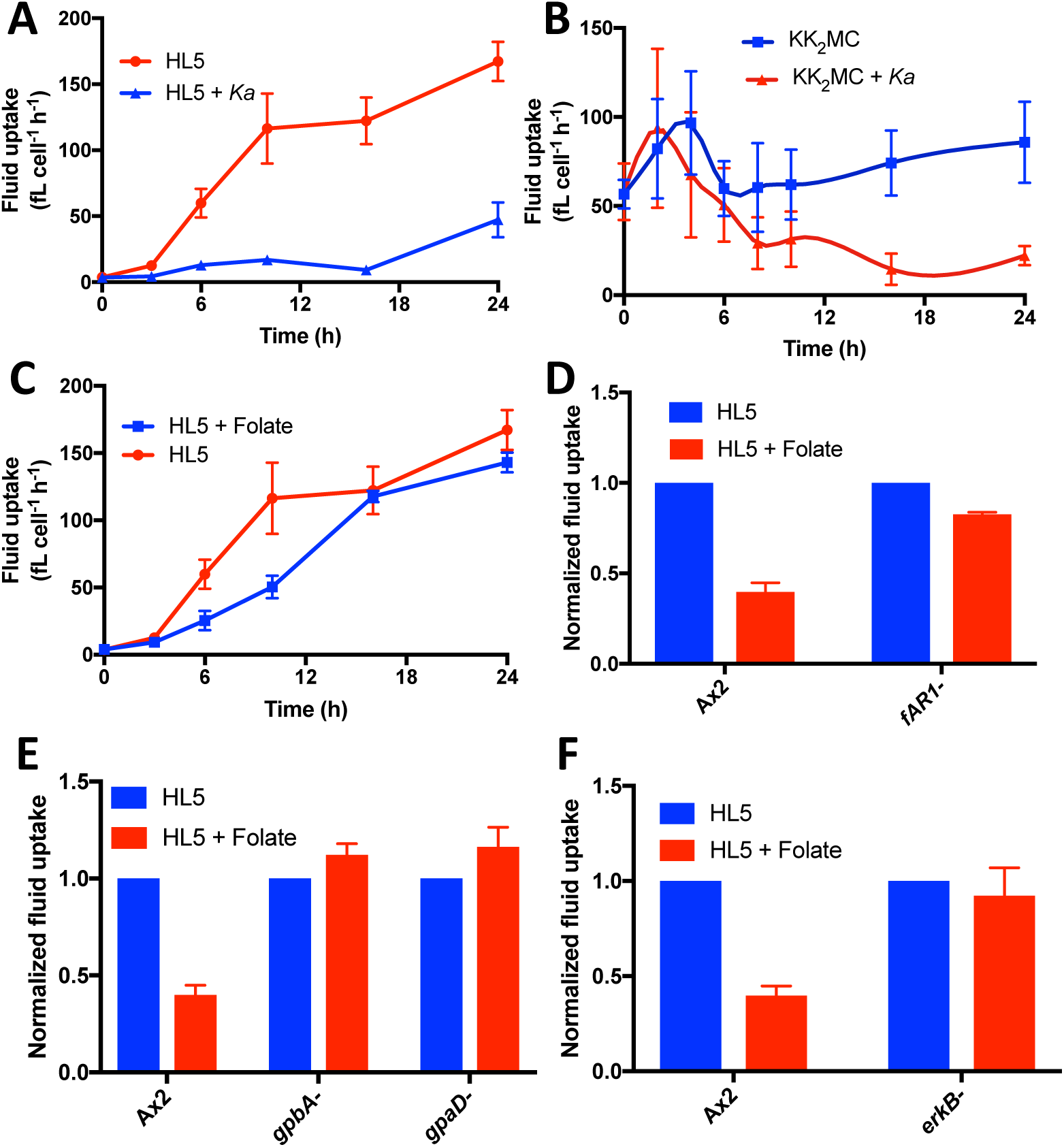
Long-term regulation of macropinocytosis by bacteria and their product, folate
**A)** Bacteria inhibit the upregulation of macropinocytosis by cells transferred to HL5 medium. Ax2 cells transferred from bacteria (low macropinocytosis) to HL5, upregulate macropinocytosis, but this is blocked by addition of *Ka* bacteria (2 OD_600 nm_) to the HL5 (n=6). **B)** Bacteria induce downregulation of macropinocytosis by cells taken from HL5 medium. Ax2 cells transferred from HL5 medium (high macropinocytosis) to KK2MC buffer maintain their rate of macropinocytosis, but the addition of 2 OD_600 nm_ *Ka* bacteria induces downregulation (n=6). **C)** Folate delays the upregulation of macropinocytosis by cells transferred to HL5 medium. Ax2 cells transferred from bacteria (low macropinocytosis) to HL5 medium, upregulate macropinocytosis, but this is delayed by 500 μM folate (n=6). **D)** The folate receptor (fAR1) mediates the inhibitory effect of folate on macropinocytosis upregulation. Wild-type Ax2 cells and a null mutant for the folate receptor (fAR1-) were transferred from bacteria (low macropinocytosis) to HL5 medium with or without 500 μM folate and macropinocytosis measured after 6 hours (n=5). **E)** The heterotrimeric G-protein cognate to the folate receptor mediates the inhibitory effect of folate on macropinocytosis upregulation. Wild-type Ax2 cells and null mutants for Gα4 (*gpaD*-) and Gβ (*gpbA*-) were transferred from bacteria (low macropinocytosis) to HL5 medium with or without 500 μM folate and macropinocytosis measured after 6 hours (n=5). **F)** The MAP-kinase, ErkB, a downstream effector of the folate receptor mediates the inhibitory effect of folate on macropinocytosis upregulation. Wild-type Ax2 cells and null mutants for ErkB (*erkB-)* were transferred from bacteria (low macropinocytosis) to HL5 medium with or without 500 μM folate and macropinocytosis measured after 6 hours (n=3).Fluid uptake was measured by high-throughput flow cytometry. Error bars are the s.e.m.

Bacteria can be sensed through their release of folate, which is a chemoattractant for *Dictyostelium* and acts through the G-protein coupled receptor fAR1 (Pan et al., 2016). We found that folate inhibits the upregulation of macropinocytosis when cells are transferred from bacteria to HL5 (Figure 5C), but has no effect when cells are transferred from HL5 to buffer (not shown). *fAR1*-cells were essentially blind to this inhibitory effect of folate (Figure 5D), as were mutants of the Gβ and Gα4 (Hadwiger and Firtel, 1992) subunits of the cognate hetero-trimeric G-protein (Figure 5E) (Hadwiger and Srinivasan, 1999) and the downstream MAP kinase, ErkB (Figure 5F) (Nichols et al., in preparation). Thus bacteria can exert some, but clearly not all, of their effects on feeding behaviour through canonical folate signalling.

### Developmental regulation of macropinocytosis

Development in *Dictyostelium* is triggered by starvation and over the first 8-10 hours the cells aggregate by chemotaxis to cyclic-AMP. Macropinocytosis is downregulated during this period (Maeda, 1983, Katoh et al., 2007), and it was therefore surprising that macropinocytosis continues at a high rate for at least 24 hours in cells starved under buffer (Figure 5B). However, compared to standard developmental conditions, these cells were starved at low density, likely causing attenuation of developmental signalling. This suggests that the downregulation of macropinocytosis during development requires a developmental signal in addition to starvation.

Figure 6A confirms that macropinocytosis is strongly downregulated by starving Ax2 cells (previously grown in HL5) at high density in shaking suspension and pulsed with cyclic-AMP to mimic developmental signalling. By 5 hours of development, fluid uptake is negligible. Similar results were obtained when developing cells on non-nutrient agar (not shown). Similarly if the cell density in 96-well plates is increased from 5000 to 50,000 cells per well, the cells can be seen to aggregate (not shown) and downregulate macropinocytosis (Figure 6B).

**Figure 6.**
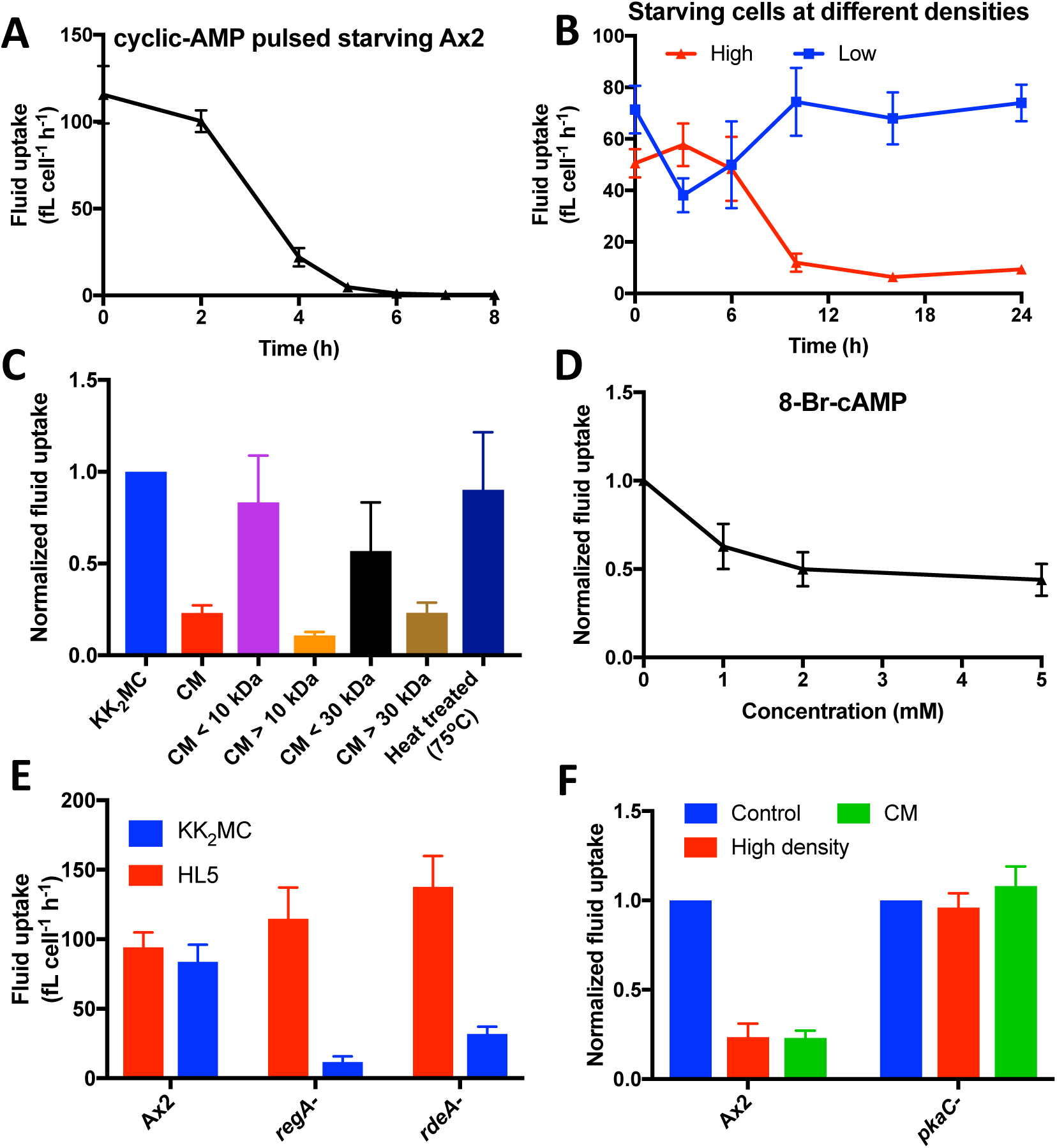
Macropinocytosis is downregulated by developmental signalling that likely acts through PKA. **A)** Macropinocytosis is downregulated during development. Ax2 cells grown in HL5 (high macropinocytosis) were washed free of nutrients and allowed to develop in standard conditions: shaken in suspension and pulsed with cyclic-AMP every 6 minutes after the first hour (n=5). **B)** Downregulation of macropinocytosis depends on the cell density. Axenically growing cells were allowed to settle at high (50,000 cells well^-1^) and low (5000 cells well^-1^) in 96-well plates, washed free of nutrient media and incubated in buffer for the indicated times before fluid uptake was determined as described in the materials and methods (n=6). **C)** Downregulation of cells at low density is induced by conditioned medium. Conditioned KK2MC (CM) prepared by shaking starving cells at high density for 8 hours was tested for its ability to induce downregulation of macropinocytosis by Ax2 cells. The CM was both size fractionated and heat-treated at 75^o^C for 30 min to further investigate the properties of the secreted product responsible for downregulation induction (n=3). **D)** Downregulation of cells at low density is induced by 8-Br-cAMP, which activates PKA, when incubated with the cells for 24 hours (n=5). **E)** Mutations giving elevated intra-cellular cyclic-AMP levels bypass the need for developmental signalling to downregulate macropinocytosis. *regA*- and *rdeA*-cells have elevated intracellular cyclic-AMP due to reduced breakdown, and downregulate macropinocytosis when incubated in KK2MC rather than HL5 for 24 hours, unlike Ax2 (n=5). **F)** Macropinocytosis is not downregulated in mutant lacking PKA activity (*pkaC*-), even when incubated in CM or at high density in buffer (n=6). Error bars show the s.e.m.

We tested the effects of known developmental signals on macropinocytosis using cells starving at low cell density. As shown in table S3, the developmental signals cyclic-AMP, ATP (Ludlow et al., 2008, Traynor and Kay, 2017), adenosine and the polyketides DIF-1, DIF-2 and MPBD (Morris et al., 1987, Morris et al., 1988, Saito et al., 2006) were without effect, as was the high cell density signal, polyphosphate (Suess and Gomer, 2016). However, conditioned medium (CM) prepared by shaking starving cells at high density for 8 hours was effective, with the active component(s) being heat-labile and retained by a 30 kDa cut-off membrane and therefore likely to be protein(s) (Figure 6C). Most likely this signal is one of the known proteins controlling early developmental events in *Dictyostelium*, but unfortunately these were unavailable for testing.

To gain insight into how developmental signals suppress macropinocytosis, we examined possible signal transduction routes, focussing on cyclic-AMP dependent protein kinase (PKA), which is a crucial mediator of both early and late events in development (Mann and Firtel, 1991, Harwood et al., 1992, Kay, 1989). PKA can be directly activated using the membrane-permeable analogue of cyclic-AMP, 8-bromo-cyclic-AMP (8-Br-cAMP) and we found that this, unlike cyclic-AMP, causes up to a 50% downregulation of macropinocytosis in starving, low-density cells (Figure 6D). High concentrations are required, but these are comparable to those used previously (Kay, 1989).

The involvement of PKA is strongly supported by mutants with elevated intracellular cyclic-AMP levels, due to defective breakdown. The hybrid cyclic-AMP phosphodiesterase RegA is activated by a His/Asp phospho-relay in which RdeA is the essential phosphate carrier protein (Shaulsky et al., 1998, Thomason et al., 1999, Thomason et al., 1998, Chang et al., 1998). Elimination of either protein results in strong downregulation of macropinocytosis in starving cells at low density (Figure 6E).

Conversely eliminating PKA activity by mutation of the catalytic subunit (*pkaC*-cells; (Primpke et al., 2000)) results in cells where macropinocytosis remains high for at least 24 hours after starvation, even if they are at high cell density or treated with CM (Figure 6F). Combined, these results strongly argue that macropinocytosis is downregulated in starving cells by PKA activation. These results are also relevant to the interpretation of recent work (Scavello et al., 2017) showing that *pkaC*-cells have a strong defect in chemotaxis to cyclic-AMP (see Discussion).

## Discussion

To facilitate work on macropinocytosis in *Dictyostelium*, we have developed a high-throughput assay to measure fluid, membrane or particle uptake with single cell resolution. While this is by no means the first use of flow cytometry for these purposes (Bacon et al., 1994, Pan et al., 2016, King et al., 2013), the high-throughput nature of the assay provides a distinct advantage over previous techniques. A screen of inhibitors provides new tools for acute inhibition of macropinocytosis and further supports the involvement of PI3K, Rac, WASP, formins and the Arp2/3 complex expected from genetic and subcellular localization studies (Hoeller et al., 2013, Langridge and Kay, 2007, Dumontier et al., 2000, Veltman et al., 2016, Junemann et al., 2016).

Macropinocytosis in *Dictyostelium* occurs at a high rate in conditions where the cells are able to proliferate in liquid medium. However it is under physiological control with cells able to slowly transition between high and low macropinocytic states according to whether bacteria or soluble nutrients are available. In these transitions the frequency of macropinosome formation is altered: in axenic cells, where the active Ras patches are unconstrained by NF1, macropinosome size is additionally increased. Wild-type cells with an intact NF1 gene also transition between low and high macropinocytic states according to the nutrients available, showing that this regulation is not just a feature of axenic strains (this work, Maeda, 1983). The presence of a high macropinocytic state in wild-type cells suggests there are ecological circumstances where macropinocytosis is used for feeding, though these are yet to be defined.

Ax2 cells in the low macropinocytic state can sense bacteria through their secretion of folic acid, inhibiting macropinocytic upregulation accordingly. However, due to the relatively modest effects of folate, and the fact that it does not induce downregulation of macropinocytosis, it seems certain that other sensory pathways also play a prominent role. It has recently been reported that certain bacteria secrete cyclic-AMP, which functions as a chemoattractant for vegetative *Dictyostelium* (Meena and Kimmel, 2017), however cyclic-AMP did not affect macropinocytosis up- or downregulation.

We find that four nutrients from defined medium are largely responsible for inducing macropinocytosis upregulation in Ax2 cells: arginine, glutamate, lysine and a metabolisable sugar. None of the other amino acids appears effective individually, and even in combination they only have a modest effect. Arginine and lysine are essential amino acids, but glutamate is not (Marin, 1976, Franke and Kessin, 1977). *Dictyostelium* has several receptors similar to metabotropic glutamate receptors (Taniura et al., 2006, Fey et al., 2013), but it seems likely that the major route for nutrient sensing is intracellular, with nutrients delivered by macropinocytosis.

In mammalian cells, free amino acids obtained by macropinocytosis are sensed through mTORC1 (Yoshida et al., 2015), which is recruited to the lysosome by Rag proteins and activated (Sancak et al., 2010). Activation of mTORC1 inhibits autophagy. *Dictyostelium* autophagy can be induced within minutes by removal of arginine and lysine (King et al., 2011), the same amino acids that induce macropinocytosis upregulation. This argues that TORC1 is activated by arginine and lysine in *Dictyostelium* to both upregulate macropinocytosis and inhibit autophagy. Though the TORC1-specific inhibitor rapamycin does not induce *Dictyostelium* autophagy (Dominguez-Martin et al., 2017), other mTor inhibitors are more effective (Cardenal-Munoz et al., 2017), suggesting that rapamycin is a relatively poor inhibitor for some *Dictyostelium* TORC1 processes, as has been suggested in mammalian cells (Thoreen and Sabatini, 2009). The more potent, but less specific, inhibitor torin 1 inhibits both macropinocytosis and macropinocytosis upregulation, although this is likely to be at least partially due to TORC2 inhibition.

As only metabolizable sugars induce upregulation of macropinocytosis, it is probable that the sensing of these is through a general metabolic readout, such as the presence of ATP, which is produced during glycolysis. Increased levels of AMP and ADP, as in nutrient poor conditions (such as without sugar), activate AMP-kinase. Overexpression of a constitutively active AMP-kinase α subunit in *Dictyostelium* inhibits growth but does not affect macropinocytosis (Bokko et al., 2007), similar to what we observe in low-density starvation conditions. It may therefore be the case that in the absence of a sugar source AMP-kinase is activated, which could prevent full macropinocytosis upregulation.

Our results show that the cessation of macropinocytosis during early development requires a developmental signal that most likely acts through PKA. Macropinocytosis does not cease immediately when cells are starved, but decreases over several hours and so may occur at reduced levels in cells that are used for studying chemotaxis to cyclic-AMP, particularly in mutants with a defect in early development. This can be a confounding influence on studies of chemotaxis, since macropinocytosis uses the same actin machinery as pseudopods and thus impairs chemotaxis (Veltman, 2015). In particular, we found that macropinocytosis continues at a high rate in mutants of the PKA catalytic subunit, possibly accounting for the strong chemotactic defect of these strains (Scavello et al., 2017). This could also be a confounding issue for some other strains with early developmental defects (Khosla et al., 2005, Wu et al., 1995, Rodriguez et al., 2008, Lee et al., 2005).

Many of the molecular components required for macropinocytosis in *Dictyostelium* are the same as those in mammalian cells: actin, Arp2/3, PI3K, SCAR/WAVE, WASP, Rac and Ras proteins. This conservation of the core components suggests that macropinocytosis may have first arisen in simple protists as a way of feeding in the absence of bacterial prey. In mammalian cells there are additional levels of regulation, some of which are cell-type specific (such as the calcium requirement in immune cells) and others that are more generic (such as growth factor stimulated macropinocytosis). We believe that *Dictyostelium* is an excellent model organism for establishing the core, conserved elements of macropinocytosis and their function.

## Materials and Methods

### Cell culture and materials

Cells were cultivated at 22°C. HL5, SIH, variants of SIH and SM media were from Formedium. Unless otherwise specified, cells were grown on *Klebsiella aerogenes* (*Ka*) lawns on SM plates and harvested for experiments from the feeding front, washing three times with KK_2_ (16.6 mM KH2PO4, 3.8 mM K2HPO4, pH 6.1) by centrifugation (280 *g*, 3 min) to remove the bacteria. Cells were also grown in tissue culture plates with *Ka* as a food source. In this case *Ka* was added to KK_2_MC (KK_2_ + 2 mM MgSO_4_, 100 μM CaCl_2_) to 2 OD_600 nm_ from a 100 OD_600 nm_ stock (these bacteria were grown overnight in 2xTY, pelleted by centrifugation and washed twice in KK_2_).

Cells were grown axenically in HL5 in conical flasks with shaking at 180 rpm. Media derived from SIH, including SUM, were made in KK_2_MC pH 6.5.Conditioned medium was made by washing axenically grown Ax2 cells free of HL5, resuspending them to 1x10^7^ cells ml^-1^ in KK2MC and incubating for 8 hours, 180 rpm, before removing the cells by centrifugation (2400 *g*, 10 min). Strains are listed in Table S4.

For transformation, cells were harvested from bacteria, resuspended in H40 buffer (40 mM Hepes, 1 mM MgCl_2_, pH 7.0), mixed with 500 ng vector for a PIP3 reporter (PkgE-PH mCherry), electroporated in ice-cold 2 mm cuvettes (Novagen) using a square wave protocol (2x 350 volts, 8 ms apart), then transferred to 2 ml KK_2_MC + *Ka* in a 6 well plate to recover for 5 hours, before G418 selection was added to 10 μg ml^-1^.

Chemicals were from Sigma unless otherwise indicated. Polyphosphate was obtained from both Spectrum and Merck.

### Uptake measurements by fluorimetry

Based on (Rivero and Maniak, 2006): cells at 1x10^7^ ml^-1^ were shaken at 180 rpm in HL5 with 0.5 mg ml^-1^ TRITC-dextran and at each time point triplicate 0.8 ml samples centrifugally washed once in ice-cold KK_2_ and resuspended to 1 ml. Fluorescence was measured in a fluorimeter (Perkin-Elmer LS 50 B with excitation at 544 nm, emission at 574 nm, slit width 10 nm). Background ‘0 minute’ fluorescence was subtracted and uptake volume calculated using standard curves of TRITC-dextran diluted in buffer. Cells loaded in this way were also analysed by flow cytometry (LSR_II flow cytometer, BD Biosciences) to compare the methods.

To measure yeast uptake, cells were resuspended from bacterial plates or 9 cm tissue culture dishes where they had been incubated in growth medium to 5x10^6^ ml^-1^ in KK2MC in a 5 ml conical flask shaken at 180 rpm, 22^o^C. TRITC-labelled yeast (sonicated at level 7.0 for 20 seconds on a Misonix sonicator 3000) were added to 1x10^7^ particles ml^-1^. At 0 and 60 minutes duplicate 200 μl samples were added to 20 μl of Trypan blue quench solution (2 mg ml^-1^ in 20 mM citrate, 150 mM NaCl, pH 4.5) on ice, shaken for 3 minutes at 2000 rpm, spun down and washed twice with ice-cold KK2 + 10 mM EDTA. The final pellet was resuspended to 1 ml and the fluorescence compared to a standard curve to give the number of yeast per cell.

### Uptake measurements by flow cytometry

For high-throughput assays, 50 μl of 1x105 cells ml^-1^ was inoculated into flat-bottom 96-well plates and incubated at 22oC for the indicated time (usually 24 hours). Then 50 μl of 1 mg ml^-1^ TRITC-dextran in the same medium was added. After one hour, unless otherwise stated, the medium was thrown off, and the cells washed by ‘dunk-banging’ (the plate was submerged in a container of ice-cold KK_2_, which was thrown off and the plate patted dry) before 100 μl KK2MC containing 5 mM sodium azide was added to each well to detach the cells and stop exocytosis. Cells were analysed by flow cytometry (LSR-II, BD Biosciences) using the High-Throughput-Sampling attachment, which pipetted them up and down twice, before analysing 65 μl per sample at 3 μl s^-1^. FlowJo software (https://www.flowjo.com) calculated the median (mean in the case of beads) fluorescence of cells in each well, and then the mean of triplicate wells was calculated. The mean was then taken of all biological replicates. To determine volumes taken up, the same population of cells (loaded with TRITC-dextran in suspension, as above) was analysed by both fluorimetry and flow cytometry. The LSR_II was calibrated through all subsequent experiments using FlowSet fluorospheres calibration beads (Beckman Coulter).

We also used this method to measure uptake of membrane using 10 μM FM1-43 (Invitrogen); phagocytosis of bacteria using 1x10^8^ particles ml^-1^ Texas-red *E. coli* bioparticles (Thermo Scientific); or beads of different sizes (YG-beads, Polysciences): 3 μm (2x10^7^ ml^-1^), 2.0 and 1.75 μm (5x10^7^ ml^-1^) or 1.5 μl^-1^ (1x10^8^ ml^-1^). Particles internalised per cell was calculated by comparing the internalised fluorescence with particles only samples. For time courses, the start time was staggered so that all time-points ended concurrently. When inhibitors were used acutely, they were added with the fluorescent medium to the final indicated concentration. Polyketides were synthesised as described (Morris et al., 1987, Morris et al., 1988, Saito et al., 2006).

To initiate development, axenically growing cells were washed twice, resuspended to 1x10^7^ cells ml^-1^ in KK_2_MC and shaken at 180 rpm for one hour before delivering pulses of KK_2_MC containing cyclic-AMP to give a concentration of 100 nM every 6 minutes using a Watson Marlow 505Di pump. At the indicated times 5x10^4^ cells were diluted into dextran containing KK_2_MC in 24 well plates for one hour, after which they were washed *in-situ* using ice-cold KK_2_ + 10 mM EDTA and detached with KK2MC + 5 mM sodium azide. 100 μl was transferred to duplicate wells in a 96-well plate for flow cytometry analysis.

Development on agar plates was initiated by settling 1.5 ml of washed, axenically grown cells at 2.5x10^7^ cells ml^-1^ in KK_2_MC onto fresh 1.8% KK_2_MC agar in 6 cm plates. After 15 minutes settling, the media was aspirated off, and the plates kept on wet tissues at 22°C. At the indicated times, cells were harvested, resuspended in KK_2_MC and 1x10^5^ inoculated into KK_2_MC in a 6-well plate with 0.5 mg ml^-1^ TRITC-dextran for one hour. Cells were then washed *in*-*situ* and resuspended in KK2 + 10 mM EDTA before analysis by low-throughput flow cytometry. The zero hour time-point was of cells taken immediately after washing.

### Macropinosome formation rate and diameter

The rate of macropinosome formation was determined in KK_2_MC by loading cells in a 2-well microscope slide (Nunc) with 2 mg ml^-1^ FITC-dextran for 1 minute, then washing and fixing with 4% paraformaldehyde for 20 minutes. Fixed cells were washed 5 times and stored in PBS (pH 5.0) at 4^o^C for imaging. Z-stacks with 0.1 μm steps were taken using a Zeiss 700 series microscope with 2x averaging to reduce noise. Maximum intensity projections were made using FIJI and FITC-positive endosomes counted by eye. The mean of at least 8 cells on a given day was taken as one data-point.

To measure macropinosome diameter at closure, cells in KK2MC expressing a PIP3 reporter (PkgE-PH mCherry) were filmed in their central section at 1 frame per second for 5 minutes on a Zeiss 700 series microscope. The maximum diameter of macropinosomes at closure was measured using the FIJI measure tool. Note that this method will underestimate the diameter of macropinosomes not lying fully within the optical section.

## Acknowledgements

We thank the rest of the Kay lab for their assistance in moulding this project, particularly Peggy Paschke. Clelia Amato and Robert Insall (Beatson Institute, Glasgow) alerted us to the Rac inhibitor. Jason King (Sheffield University) provided valuable feedback on the macropinosome formation experiments. Miao Pan and Tian Jin (NIAID, Bethesda) kindly sent us the *fAR1-* strain. The MRC-LMB flow cytometry facility maintained the flow cytometers and provided technical support.

## Competing Interests

The authors declare that there are no competing interests.

## Author Contributions

Both authors designed the experiments, which were carried out by Thomas Williams.

## Funding

We thank the Medical Research Council UK for core funding (U105115237 to RRK).

## Data availability

N/A.

## References

Aguado-Velasco, C. & Bretscher, M. S. 1999. Circulation of the plasma membrane in Dictyostelium. Mol. Biol. Cell, 10, 4419-4427.

Ashworth, J. M. & Watts, D. J. 1970. Metabolism of the cellular *slime mould Dictyostelium discoideum grown* in axenic culture. Biochem. J., 119, 175-182.

Bacon, R. A., Cohen, C. J., Lewin, D. A. & Mellman, I. 1994. *Dictyostelium discoideum* mutants with temperature-sensitive defects in endocytosis. J. Cell Biol., 127, 387-399.

Bloomfield, G., Traynor, D., Sander, S. P., Veltman, D. M., Pachebat, J. A. & Kay, R. R. 2015. Neurofibromin controls macropinocytosis and phagocytosis in Dictyostelium. Elife, 4, p. e04940.

Bokko, P. B., Francione, L., Bandala-Sanchez, E., Ahmed, A. U., Annesley, S. J., Huang, X., Khurana, T., Kimmel, A. R. & Fisher, P. R. 2007. Diverse cytopathologies in mitochondrial disease are caused by amp-activated protein kinase signaling. Mol. Biol. Cell, 18, 1874-1886.

Buckley, C. M. & King, J. S. 2017. Drinking problems: mechanisms of macropinosome formation and maturation. FEBSJ., 284, 3778-3790.

Canton, J., Schlam, D., Breuer, C., Gutschow, M., Glogauer, M. & Grinstein, S. 2016. Calcium-sensing receptors signal constitutive macropinocytosis and facilitate the uptake of NOD2 ligands in macrophages. Nat. Commun., 7, 11284.

Cardenal-Munoz, E., Arafah, S., Lopez-Jimenez, A. T., Kicka, S., Falaise, A., Bach, F., Schaad, O., King, J. S., Hagedorn, M. & Soldati, T. 2017. *Mycobacterium marinum* antagonistically induces an autophagic response while repressing the autophagic flux in a TORC1- and ESX-1-dependent manner. PLoS Pathog., 13, e1006344.

Chang, W. T., Thomason, P. A., Gross, J. D. & Newell, P. C. 1998. Evidence that the RdeA protein is a component of a multistep phosphorelay modulating rate of development in *Dictyostelium*. EMBO J., 17, 2809-2816.

Clark, J., Kay, R. R., Kielkowska, A., Niewczas, I., Fets, L., Oxley, D., Stephens, L. R. & Hawkins, P. T. 2014. *Dictyostelium* uses ether-linked inositol phospholipids for intracellular signalling. EMBO J., 33, 2188-200.

Commisso, C., Davidson, S. M., Soydaner-Azeloglu, R. G., Parker, S. J., Kamphorst, J. J., Hackett, S., Grabocka, E., Nofal, M., Drebin, J. A., Thompson, C. B., Rabinowitz, J. D., Metallo, C. M., Vander Heiden, M. G. & Bar-Sagi, D. 2013. Macropinocytosis of protein is an amino acid supply route in Ras-transformed cells. Nature, 497, 633-7.

Dominguez-Martin, E., Cardenal-Munoz, E., King, J. S., Soldati, T., Coria, R. & Escalante, R. 2017. Methods to Monitor and Quantify Autophagy in the Social Amoeba *Dictyostelium discoideum*. Cells, 6.

Dumontier, M., Hocht, P., Mintert, U. & Faix, J. 2000. Rac1 GTPases control filopodia formation, cell motility, endocytosis, cytokinesis and development in *Dictyostelium*. J. Cell Sci., 113, 2253-2265.

Falcon, B., Cavallini, A., Angers, R., Glover, S., Murray, T. K., Barnham, L., Jackson, S., O'Neill, M. J., Isaacs, A. M., Hutton, M. L., Szekeres, P. G., Goedert, M. & Bose, S. 2015. Conformation determines the seeding potencies of native and recombinant Tau aggregates. J. Biol. Chem., 290, 1049-65.

Fares, H. & Greenwald, I. 2001. Genetic analysis of endocytosis in *Caenorhabditis elegans:* coelomocyte uptake defective mutants. Genetics, 159, 133-45.

Fevrier, B., Vilette, D., Archer, F., Loew, D., Faigle, W., Vidal, M., Laude, H. & Raposo, G. 2004. Cells release prions in association with exosomes. Proc. Natl. Acad. Sci. USA, 101, 9683-8.

Fey, P., Dodson, R. J., Basu, S. & Chisholm, R. L. 2013. One stop shop for everything *Dictyostelium:* dictyBase and the Dicty Stock Center in 2012. Methods Mol. Biol., 983, 59-92.

Franke, J. & Kessin, R. 1977. A defined minimal medium for axenic strains of *Dictyostelium discoideum*. Proc. Natl. Acad. Sci. USA, 74, 2157-2161.

Glynn, P. J. & Clarke, K. R. 1984. An investigation of adhesion and detachment in slime mould amoebae using columns of hydrophobic beads. Exp. Cell Res., 152, 117-126.

Hacker, U., Albrecht, R. & Maniak, M. 1997. Fluid-phase uptake by macropinocytosis in *Dictyostelium*. J. Cell Sci., 110, 105-112.

Hadwiger, J. A. & Firtel, R. A. 1992. Analysis of Galpha4, a G-protein subunit required for multicellular development in *Dictyostelium*. Genes Devel., 6, 38-49.

Hadwiger, J. A. & Srinivasan, J. 1999. Folic acid stimulation of the Galpha4 G protein-mediated signal transduction pathway inhibits anterior prestalk cell development in *Dictyostelium*. Differentiation, 64, 195-204.

Hardt, W. D., Chen, L. M., Schuebel, K. E., Bustelo, X. R. & Galan, J. E. 1998. *S. typhimurium* encodes an activator of Rho GTPases that induces membrane ruffling and nuclear responses in host cells. Cell, 93, 815-26.

Harwood, A. J., Hopper, N. A., Simon, M. N., Bouzid, S., Veron, M. Williams, J. G. 1992. Multiple roles for cAMP-dependent protein kinase during *Dictyostelium* development. Dev. Biol., 149, 90-99.

Hoeller, O., Bolourani, P., Clark, J., Stephens, L. R., Hawkins, P. T., Weiner, O. D., Weeks, G. & Kay, R. R. 2013. Two distinct functions for PI3-kinases in macropinocytosis. J. Cell Sci., 126, 4296-307.

Junemann, A., Filic, V., Winterhoff, M., Nordholz, B., Litschko, C., Schwellenbach, H., Stephan, T., Weber, I. & Faix, J. 2016. A Diaphanous-related formin links Ras signaling directly to actin assembly in macropinocytosis and phagocytosis. Proc. Natl. Acad. Sci. USA, 113, E7464-E7473.

Katoh, M., Chen, G., Roberge, E., Shaulsky, G. & Kuspa, A. 2007. Developmental commitment in *Dictyostelium discoideum*. Eukaryot. Cell, 6, 2038-2045.

Kay, R. R. 1989. Evidence that elevated intracellular cyclic AMP triggers spore maturation in *Dictyostelium*. Development, 105, 753-759.

Kayman, S. C. & Clarke, M. 1983. Relationship between axenic growth of *Dictyostelium discoideum* strains and their track morphology on substrates coated with gold particles. J. Cell Biol., 97, 1001-1010.

Khosla, M., Splegelman, G. B. & Weeks, G. 2005. The effect of the disruption of a gene encoding a PI4 kinase on the developmental defect exhibited by *Dictyostelium* rasC-cells. Dev. Biol., 284, 412-420.

King, J. S., Gueho, A., Hagedorn, M., Gopaldass, N., Leuba, F., Soldati, T. & Insall, R. H. 2013. WASH is required for lysosomal recycling and efficient autophagic and phagocytic digestion. Mol. Biol. Cell, 24, 2714-26.

King, J. S., Veltman, D. M. & Insall, R. H. 2011. The induction of autophagy by mechanical stress. Autophagy, 7, 1490-1499.

Koivusalo, M., Welch, C., Hayashi, H., Scott, C. C., Kim, M., Alexander, T., Touret, N., Hahn, K. M. & Grinstein, S. 2010. Amiloride inhibits macropinocytosis by lowering submembranous pH and preventing Rac1 and Cdc42 signaling. J. Cell Biol., 188, 547-63.

Langridge, P. D. & Kay, R. R. 2007. Mutants in the *Dictyostelium* Arp2/3 complex and chemoattractant-induced actin polymerization. Exp. Cell Res., 313, 2563-2574.

Lee, S., Comer, F. I., Sasaki, A., Mcleod, I. X., Duong, Y., Okumura, K., Yates, J. R., Parent, C. A. & Firtel, R. A. 2005. TOR complex 2 integrates cell movement during chemotaxis and signal relay in *Dictyostelium*. Mol. Biol. Cell, 16, 4572-4583.

Lewis, W. H. 1931. Pinocytosis. Johns Hopkins Hosp. Bull., 49, 17-27.

Lewis, W. H. 1937. Pinocytosis by malignant cells. Cancer Res., 29, 666-679.

Ludlow, M. J., Traynor, D., Fisher, P. R. & Ennion, S. J. 2008. Purinergic-mediated Ca2+ influx in Dictyostelium discoideum. Cell Calcium, 44, 567-579.

Lukyanenko, V., Malyukova, I., Hubbard, A., Delannoy, M., Boedeker, E., Zhu, C., Cebotaru, L. & Kovbasnjuk, O. 2011. Enterohemorrhagic *Escherichia coli* infection stimulates Shiga toxin 1 macropinocytosis and transcytosis across intestinal epithelial cells. Am. J. Physiol. Cell Physiol., 301, C1140-9.

Maeda, Y. 1983. Axenic growth of *Dictyostelium discoideum* wild-type NC-4 cells and its relation to endocytotic ability. J. Gen. Microbiol., 129, 2467-2473.

Maeda, Y. 1988. Changes of endocytotic activities during the cell cycle of *Dictyostelium* cells. Devel. Growth Differ., 30, 15-24.

Maeda, Y. & Kawamoto, T. 1986. Pinocytosis in *Dictyostelium discoideum* cells. A possible implication of cytoskeletal actin for pinocytotic activity. Exp. Cell Res., 164, 516-526.

Magzoub, M., Sandgren, S., Lundberg, P., Oglecka, K., Lilja, J., Wittrup, A., Goran Eriksson, L. E., Langel, U., Belting, M. & Graslund, A. 2006. N-terminal peptides from unprocessed prion proteins enter cells by macropinocytosis. Biochem. Biophys. Res. Commun., 348, 379-85.

Mann, S. K. O. & Firtel, R. A. 1991. A developmentally regulated, putative serine/threonine protein kinase is essential for development in *Dictyostelium*. Mech. Devel., 35, 89-101.

Marechal, V., Prevost, M. C., Petit, C., Perret, E., Heard, J. M. & Schwartz, O. 2001. Human immunodeficiency virus type 1 entry into macrophages mediated by macropinocytosis. J. Virol., 75, 11166-77.

Marin, F. T. 1976. Regulation of development in *Dictyostelium discoideum:* I. Initiation of the growth to developmental transition by amino acid starvation. Dev. Biol., 48, 110-117.

Meena, N. P. & Kimmel, A. R. 2017. Chemotactic network responses to live bacteria show independence of phagocytosis from chemoreceptor sensing. Elife, 6.

Morris, H. R., Masento, M. S., Taylor, G. W., Jermyn, K. A. & Kay, R. R. 1988. Structure elucidation of two differentiation inducing factors (DIF-2 and DIF-3) from the cellular slime mould Dictyostelium discoideum. Biochem. J, 249, 903-906.

Morris, H. R., Taylor, G. W., Masento, M. S., Jermyn, K. A. & Kay, R. R. 1987. Chemical structure of the morphogen differentiation inducing factor from Dictyostelium discoideum. Nature, 328, 811-814.

Munch, C., O'Brien, J. & Bertolotti, A. 2011. Prion-like propagation of mutant superoxide dismutase-1 misfolding in neuronal cells. Proc. Natl. Acad. Sci. USA, 108, 3548-53.

Nanbo, A., Imai, M., Watanabe, S., Noda, T., Takahashi, K., Neumann, G., Halfmann, P. & Kawaoka, Y. 2010. *Ebolavirus* is internalized into host cells via macropinocytosis in a viral glycoprotein-dependent manner. PLoS Pathog., 6, e1001121.

Norbury, C. C., Hewlett, L. J., Prescott, A. R., Shastri, N. & Watts, C. 1995. Class I MHC presentation of exogenous soluble antigen via macropinocytosis in bone marrow macrophages. Immunity, 3, 783-91.

Novak, K. D., Peterson, M. D., Reedy, M. C. & Titus, M. A. 1995. *Dictyostelium* myosin I double mutants exhibit conditional defects in pinocytosis. J. Cell Biol., 131, 1205-1221.

Pan, M., Xu, X., Chen, Y. & Jin, T. 2016. Identification of a Chemoattractant G-Protein-Coupled Receptor for Folic Acid that Controls Both Chemotaxis and Phagocytosis. Dev. Cell, 36, 428-39.

Parent, C. A., Blacklock, B. J., Froelich, W. M., Murphy, D. B. & Devreotes, P. N. 1998. G Protein signaling events are activated at the leading edge of chemotactic cells. Cell, 95, 81-91.

Patel, H. & Barber, D. L. 2005. A developmentally regulated Na-H exchanger in *Dictyostelium discoideum* is necessary for cell polarity during chemotaxis. J. Cell Biol., 169, 321-329.

Pramanik, M. K., Iijima, M., Iwadate, Y. & Yumura, S. 2009. PTEN is a mechanosensing signal transducer for myosin II localization in *Dictyostelium* cells. Genes to Cells, 14, 821-834.

Primpke, G., Iassonidou, V., Nellen, W. & Wetterauer, B. 2000. Role of cAMP-dependent protein kinase during growth and early development of *Dictyostelium discoideum*. Dev. Biol., 221, 101-111.

Rivero, F. & Maniak, M. 2006. Quantitative and microscopic methods for studying the endocytic pathway. Methods Mol. Biol., 346, 423-438.

Rodriguez, M., Kim, B., Lee, N. S., Veeranki, S. & Kim, L. 2008. MPL1, a novel phosphatase with leucine-rich repeats, is essential for proper ERK2 phosphorylation and cell motility. Euk. Cell., 7, 958-966.

Rosel, D., Khurana, T., Majithia, A., Huang, X., Bhandari, R. & Kimmel, A. R. 2012. TOR complex 2 (TORC2) in *Dictyostelium* suppresses phagocytic nutrient capture independently of TORC1-mediated nutrient sensing. J. Cell Sci., 125, 37-48.

Saito, T., Taylor, G. W., Yang, J. C., Neuhaus, D., Stetsenko, D., Kato, A. & Kay, R. R. 2006. Identification of new differentiation inducing factors from *Dictyostelium discoideum*. Biochim. Biophys. Acta, 1760, 754-761.

Sallusto, F., Cella, M., Danieli, C. & Lanzavecchia, A. 1995. Dendritic cells use macropinocytosis and the mannose receptor to concentrate macromolecules in the major histocompatibility complex class II compartment: downregulation by cytokines and bacterial products. J. Exp. Med., 182, 389-400.

Sancak, Y., Bar-Peled, L., Zoncu, R., Markhard, A. L., Nada, S. & Sabatini, D. M. 2010. Ragulator-Rag complex targets mTORC1 to the lysosomal surface and is necessary for its activation by amino acids. Cell, 141, 290-303.

Scavello, M., Petlick, A. R., Ramesh, R., Thompson, V. F., Lotfi, P. & Charest, P. G. 2017. Protein kinase A regulates the Ras, Rap1 and TORC2 pathways in response to the chemoattractant cAMP in *Dictyostelium*. J. Cell Sci., 130, 1545-1558.

Shaulsky, G., Fuller, D. & Loomis, W. F. 1998. A cAMP-phosphodiesterase controls PKA-dependent differentiation. Development, 125, 691-699.

Shu, S., Liu, X. & Korn, E. D. 2005. Blebbistatin and blebbistatin-inactivated myosin II inhibit myosin II-independent processes in *Dictyostelium*. Proc. Natl. Acad. Sci. USA, 102, 1472-1477.

Shutes, A., Onesto, C., Picard, V., Leblond, B., Schweighoffer, F. & Der, C. J. 2007. Specificity and mechanism of action of EHT 1864, a novel small molecule inhibitor of Rac family small GTPases. J. Biol. Chem., 282, 35666-78.

Suess, P. M. & Gomer, R. H. 2016. Extracellular Polyphosphate Inhibits Proliferation in an Autocrine Negative Feedback Loop in *Dictyostelium discoideum*. J. Biol. Chem., 291, 20260-9.

Taniura, H., Sanada, N., Kuramoto, N. & Yoneda, Y. 2006. A metabotropic glutamate receptor family gene in *Dictyostelium discoideum*. J. Biol. Chem., 281, 12336-12343.

Thilo, L. & Vogel, G. 1980. Kinetics of membrane internalization and recycling during pinocytosis in *Dictyostelium discoideum*. Proc. Natl. Acad. Sci. USA, 77, 1015-1019.

Thomason, P. A., Traynor, D., Cavet, G., Chang, W.-T., Harwood, A. J. & Kay, R. R. 1998. An intersection of the cAMP/PKA and two-component signal transduction systems in *Dictyostelium*. EMBO J., 17, 2838-2845.

Thomason, P. A., Traynor, D., Stock, J. B. & Kay, R. R. 1999. The RdeA-RegA system, a eukaryotic phospho-relay controlling cAMP breakdown. J. Biol. Chem., 274, 27379-27384.

Thoreen, C. C. & Sabatini, D. M. 2009. Rapamycin inhibits mTORC1, but not completely. Autophagy, 5, 725-6.

Traynor, D. & Kay, R. R. 2017. A polycystin-type transient receptor potential (Trp) channel that is activated by ATP. Biol Open, 6, 200-209.

Veltman, D. M. 2015. Drink or drive: competition between macropinocytosis and cell migration. Biochem. Soc. Trans., 43, 129-32.

Veltman, D. M., Williams, T. D., Bloomfield, G., Chen, B. C., Betzig, E., Insall, R. H. & Kay, R. R. 2016. A plasma membrane template for macropinocytic cups. Elife, 5, e20085.

Watts, D. J. & Ashworth, J. M. 1970. Growth of myxamoebae of the cellular slime mould *Dictyostelium discoideum* in axenic culture. Biochem. J., 119, 171-174.

Wu, L. J., Valkema, R., Van Haastert, P. J. M. & Devreotes, P. N. 1995. The G protein beta subunit is essential for multiple responses to chemoattractants in *Dictyostelium*. J. Cell Biol., 129, 1667-1675.

Yoshida, S., Pacitto, R., Yao, Y., Inoki, K. & Swanson, J. A. 2015. Growth factor signaling to mTORC1 by amino acid-laden macropinosomes. J. Cell Biol., 211, 159-72.

